# The variability of song variability in wild and domesticated zebra finches *Taeniopygia guttata*

**DOI:** 10.1101/263913

**Authors:** Allison L. Lansverk, Sarah E. London, Simon C. Griffith, David F. Clayton, Christopher N. Balakrishnan

## Abstract

Birdsong is a classic example of a learned social behavior. Like many traits of interest, however, song production is also influenced by genetic factors and understanding the relative contributions of genetic and environmental influences remains a major research goal. In this study we take advantage of genetic variation among captive zebra finch populations to examine variation in a population-level song trait: song variability. We find that zebra finch populations differ in levels of song variability. Domesticated *T. g. castanotis* populations displayed higher song diversity than more recently wild-derived populations of both zebra finch subspecies *T. g. castanotis* and *T. g. guttata*, the Timor zebra finch. To determine whether these differences could have a genetic basis, we cross-fostered domesticated *T. g. castanotis* and Timor zebra finches to Bengalese finches *Lonchura striata domestica*. Following cross-fostering, domesticated *T. g. castanotis* maintained a higher level of song diversity than *T. g. guttata*. We suggest that the high song variability of domesticated zebra finches may be a consequence of reduced purifying selection acting on song traits. Intraspecific differences in the mechanisms underlying song variability therefore represent an untapped opportunity for probing the mechanisms of song learning and production.

## 1. Introduction

As a representative of the Oscine Passerines, or songbirds, zebra finches have been the subject of extensive neurobiological and behavioral research with a focus on vocal communication behavior (Marler 1970; ten Cate 2014; Jin and Clayton 1997; Dave and Margoliash 2000; Olveczky et al. 2005; Mooney 2009; London and Clayton 2008; Thompson et al. 2011; Vallentin et al. 2016). Despite its role as a model system, however, zebra finch song behavior is in some ways unusual among songbirds. For example, zebra finches are highly age-restricted or “closed” learners that don’t modify their songs after around 90 days of age (Brenowitz and Beecher 2005). Zebra finches also maintain high song structural diversity even within local populations, yet show little evidence for the regional song dialects well-known in songbirds (Lachlan et al. 2016; Marler and Tamura 1962).

As zebra finches are highly colonial birds, population-level song structural diversity may facilitate individual recognition. Exposure to novel song has been shown to cause differential behavioral, electrophysiological and gene expression responses relative to playbacks of songs to which birds have previously been exposed (reviewed in Dong and Clayton 2009), and this recognition learning must be dependent in part on salient structural variation in song. Juvenile zebra finches regularly produce novel syllables that are distinctive from those of their tutors (Jones et al. 1996; Houx and ten Cate 1999; Holveck et al. 2008). Lachlan et al. (2016) suggested that the lack of local dialects may be the result of high variability in zebra finch song.

In this study, we test for population differences in song variability itself. The majority of zebra finch research relies on domesticated populations of the *T. g. castanotis* subspecies, native to Australia. Relative to many domesticated species, the domestication of the zebra finch has been recent (~150 years) and closely-related wild populations are still available for comparison (Rogers 1979; Clayton 1989, Zann 1996). Previous studies have demonstrated significant genetic differentiation between domesticated and wild populations and a loss of genetic diversity in captive populations (Forstmeier et al. 2007). There is also evidence for human-mediated selection on zebra finches including selection for larger body size (Zann 1996, Forstmeier et al. 2007). Unlike many lab models, however, captive zebra finch populations are often kept as more natural, free-mating, outbred colonies. Thus, despite a loss of genetic diversity there is no evidence of severe genetic bottlenecks (Forstmeier et al 2007).

Timor zebra finches *T. g. guttata*, native to the Lesser Sunda islands of southeast Asia, have more recently been established in the pet trade and are also beginning to see research use (Perfito et al. 2008; Hofmeister et al 2017). The colonization of the Lesser Sunda islands by zebra finches appears to have occurred about one million years ago and was accompanied by a severe population bottleneck (Balakrishnan and Edwards 2009). Previous work has described differences between *T. g. castanotis and T. g. guttata* in song length, frequency, and the number of elements in a phrase (Clayton 1990). Divergence in the wild and during the process of domestication also have the potential to influence aspects of population song variability in zebra finches.

## 2. Materials & Methods

### (a) Study populations

We recorded songs of individuals sampled from three populations of *T. g. castanotis* (East Carolina University (ECU), n = 10), University of Chicago (n = 8) and Macquarie University (n = 7) and one population of *T. g. guttata* ((“Timor”) n = 10). We consider the ECU and Chicago populations to be domesticated in that their origins trace to long-term captive populations in the US. The ECU colony of *T. g. castanotis* was founded by five pairs of birds in 2012. These five pairs were drawn from another captive population housed at ECU (courtesy of Dr. Ken Soderstrom) which was itself founded in 2002 with birds sourced from Acadiana Aviaries (Franklin, Louisiana, USA). The Chicago population was derived from adult birds sourced from Magnolia Bird Farm (Anaheim, CA, USA). One Chicago colony was originally founded in October 2011 by a set of 40 males and 40 females. In June 2013 another colony of 50 males and 50 females was founded. Each of these two populations at Chicago bred independently until they were merged in November 2015. Birds recorded here were drawn from this merged colony.

Two others colonies, *T. g. castanotis* at Macquarie and *T. g. guttata* at ECU were more recently brought into captivity from the wild. The Macquarie University colony was founded in 2007 using approximately 100 birds taken directly from the wild in Australia, with an additional 40 birds (also taken directly from the wild) added to the population in 2010. These wild-derived birds have been kept isolated from domesticated birds and have had the opportunity to breed about every 12 months (Gilby et al. 2013). The ancestors of the Timor zebra finches used in this study were originally brought into captivity in the US in the early 1990s (efinch.com, San Jose CA, USA). The colony used here was founded by five pairs of individuals in 2009 and have been maintained at 40-50 individuals since 2012. All of the zebra finches used in this study were bred, raised and housed in large flight aviaries and thus were exposed to similar acoustic environments and a rich social environment.

### (b) Experimental tutoring

In order to compare song copying behavior between populations that differ in song variability (see below), we conducted a tutoring experiment with the *T. g. castanotis* and *T. g. guttata* housed at ECU. Juvenile zebra finches were tutored by Bengalese finches, *Lonchura striata domestica*. This approach has been used successfully used many times in experiments with zebra finches (e.g., Eales 1987a; ten Cate 1987; Clayton 1989; Takahasi et al. 2006; Campbell and Hauber 2009; Campbell and Hauber 2009; Soma 2011; Villain et al. 2015). In each case, a male zebra finch was placed with a male Bengalese finch tutor in a small cage; these cages had visual barriers between them so they could not see other tutors or cross-fostered zebra finches. These barriers also provided partial acoustic isolation. The housing room also contained flight aviaries with both zebra finch subspecies and Bengalese finches but tutoring cages were spatially separated from the aviaries to minimize social interactions between cages and aviaries. In total, five different Bengalese finch tutors were used, balanced among treatments as evenly as was feasible, with each Bengalese finch tutoring at least one individual of each subspecies.

For logistical reasons we used two different strategies for tutoring, moving eggs to foster parents (n = 3 *T. g. castanotis*, n = 2 *T. g. guttata*), and moving birds at post-hatch day 30 (p30) to tutoring cages (n = 7 *T. g. castanotis*, n = 6 *T. g. guttata*). Cross-fostered eggs were often abandoned, whereas by p30 zebra finches were feeding independently, greatly facilitating the tutoring experiments. All individuals were left with their tutors until at least day 90, after the close of the critical period for song learning and song crystallization (Immelmann 1969, Price 1979).

### (c) Song Recording & Analysis

Pairs of birds (one male and one female) were placed in a sound chamber and recorded using the activity-triggered recording in Sound Analysis Pro 2011 (SAP2011) software (Tchernichovski et al. 2000). Timor zebra finches rarely vocalized in the absence of a female so this approach was required in order to collect comparable song data. By pairing a male and female, we captured a mixture of both female-directed and undirected song (Sossinka 1980, Woolley and Doupe 2008; Lachlan et al. 2016). ECU and Timor populations were recorded from 2014 to 2016 at ECU using a Sennheiser ME22 shotgun microphone connected to a Focusrite Scarlett 2i2 USB pre-amplifier, which was in turn connected to IBM Thinkpad laptop running SAP. The wild-derived colony at Macquarie University was recorded in July 2016 at Macquarie using the same equipment. Birds at the University of Chicago were recorded during April 2017 using a Rode NT5 Small-diaphragm Cardioid Condenser Microphones connected to Focusrite Scarlett 2i2 USB Pre-amplifiers; songs were recorded via Dell OptiPlex desktop computers running SAP2011. Birds were left in the chamber until the males produced at least 100 song bouts (generally 2-3 days). This detailed characterization of individual repertoires is required for the song analysis techniques described below.

SAP2011 was also used for quantitative comparisons of song structure. The batch analysis function in SAP creates syllable tables that gives information for 14 song parameters: syllable duration, mean amplitude, mean pitch, mean frequency modulation (FM), mean squared amplitude modulation (AM^2^), mean Weiner entropy, mean pitch goodness, mean frequency, variance in pitch, variance in FM, variance in entropy, variance in pitch goodness, variance in mean frequency, and variance in AM. Following recording, syllables shorter than 20 milliseconds were filtered to remove cage noise from recordings. We used the Feature Batch function in SAP to parse the motifs into syllables by setting segmentation values for amplitude, mean frequency and continuity, which are unique to each individual.

To quantify population-level song variability we estimated Kullback-Leibler (K-L) divergence (Wu et al. 2008) using KLfromRecordingDays (Soderstrom and Alalawi 2017) in pairwise comparisons among individuals within populations (Macquarie, Chicago, ECU and Timor) and within groups of cross-fostered birds (ECU and Timor). These pairwise estimates of K-L divergence included comparisons in which each bird was used as both a “template” and a “target” to generate a matrix of pairwise comparisons. Because K-L distance is an asymmetric measurement, we took the average of both comparisons using each pair of birds to generate an estimate of song similarity between each pair of birds. We also used K-L distance to quantify similarity between cross-fostered birds and their tutors (ECU and Timor). All K-L divergence estimates include syllable duration as one of the variables, but were estimated for the remaining 13 secondary song parameters listed above.

All statistical analyses of estimated K-L distances were performed using R (R Core Team 2014). We used K-L distances as a response measure in linear models (*Imer* function in *Ime4*) with Population as a fixed factor. For analyses of population song variability we treated the individual birds used in each comparison as a random factor to accommodate repeated comparisons. For analyses of song similarity between ECU *T. g. castanotis, T. g. guttata* and their Bengalese finch tutors, we again use K-L distances as a response measure in linear models, but included the ID of Bengalese finch tutor as a random factor in the model. Because K-L distances for different song metrics were highly correlated, we used PCA to summarize distances in a single variable and used PC1 as a response measure in linear models as above. We tested for overall population differences in variability using a likelihood ratio test and then used pairwise post-hoc tests using *difflsmeans* in the *ImerTest* package.

## 3. Results

In an overall comparison of three *T. g. castantotis* and one *T. g. guttata* colonies, we found a significant effect of source population on each acoustic measurement as well as the summary principal component 1 (PC1) score (Table 1, Figure 1), which explained 78.6% of the variance in the data. Pairwise post-hoc tests revealed a significant difference between the ECU population and each of the other sampled populations in PC1 (based on all K-L distance estimates) and all individual song parameters, with ECU birds showing consistently high variability (mean K-L distance). The other domesticated population, sampled from the University of Chicago, was not significantly different from the wild-derived Macquarie population in PC1 (estimated least-squares difference in means (LSD) = −1.1, *t* = −1.25, *p* = 0.22), but did show significantly different K-L distances for individual measures FM (LSD = −0.7, *t* = −2.04, *p* = 0.05), entropy (LSD = − 1.2, *t* = −2.78, *p* = 0.009) and variance in pitch (LSD = −0.4, *t* = −2.21, *p* = 0.034). Timor zebra finches tended to show lower song variability than the three *T. guttata castanotis* populations (Table 1). In comparisons between *T. g. guttata* and the wild-derived Macquarie *T. g. castanotis* this difference was only statistically significant for amplitude modulation (LSD = 0.7, *t* = 2.27, *p* = 0.029), variance in FM (LSD = 1.1, *t* = 3.64, *p* = 9e-04), and variance in entropy (LSD = 1.0, *t* = 3.24, *p* = 0.003).

**Table 1.**
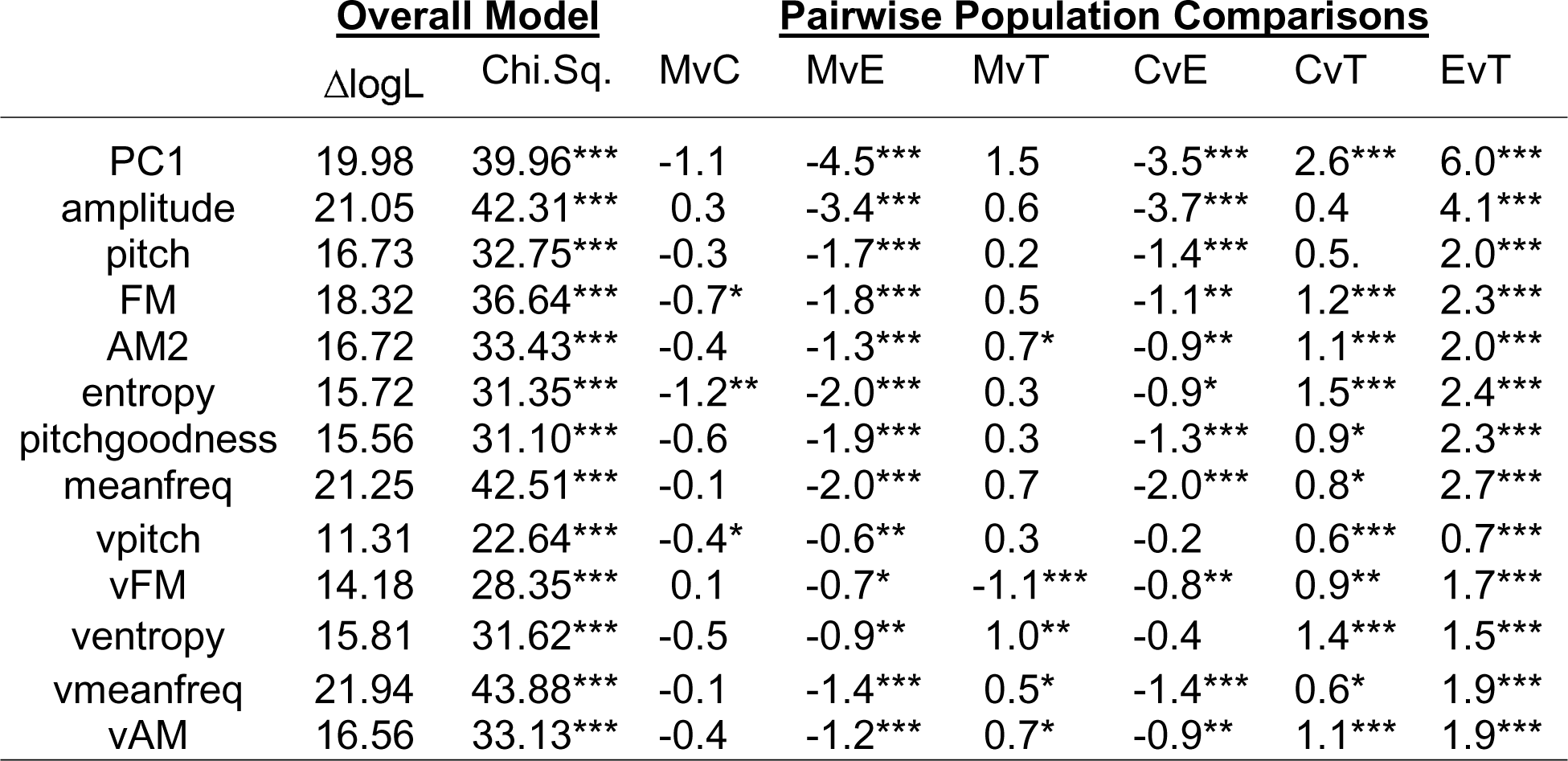
Overall test for population differences in song variability as measured for thirteen song properties and a principal component score (PC1) summarizing those measurements. A likelihood ratio test with one degree of freedom was used to compare a model with “Population” as a fixed factor with a null model. Post-hoc tests were then used to compare (M)acquarie, C(hicago), E(CU) and T(imor) finches. Presented are estimated least squares differences between populations and statistical significance based on a t-test (* p < 0.05, ** p <0.01, *** p < 0.001). Negative values indicate that the first population in the comparison had lower variability (fitted mean pairwise K-L distance) and positive values indicate that the first population had higher K-L distances between individuals.

**Figure 1.**
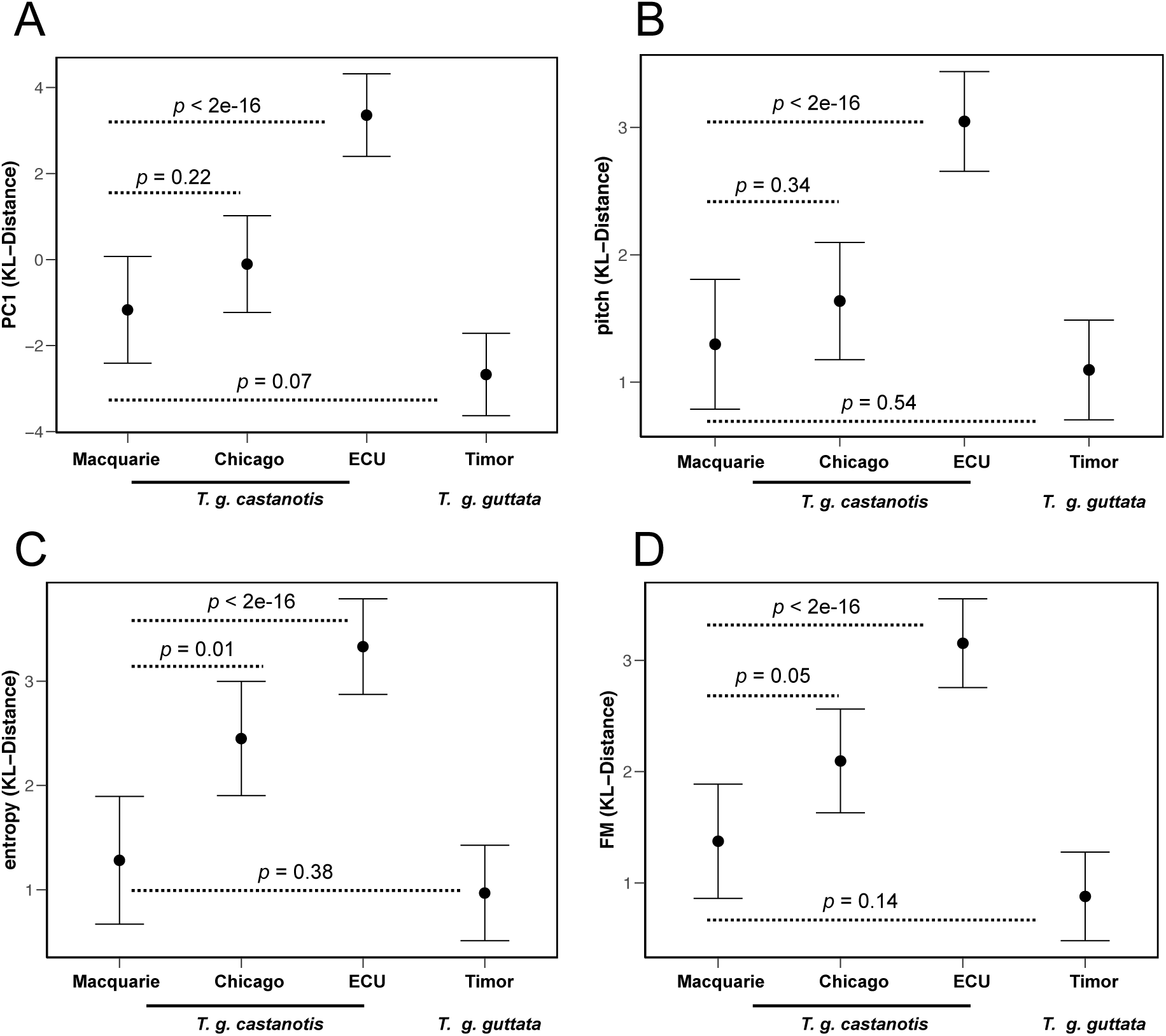
Fitted means and confidence intervals for population estimates of KL-distance for A) Principal Component 1 B) pitch, C) entropy and D) frequency modulation (FM). Higher mean K-L distance indicates higher song variability. Dashed lines highlight pairwise *post-hoc* t-tests and associated p-values between wild-derived birds from Macquarie and the other three colonies. The results of all pairwise comparisons are provided in Table 1.

Following cross-fostering there was no difference in K-L distance between *T. g. castanotis* and *T. g. guttata* housed at ECU in terms of how closely they matched their tutor songs (ΔlogL = 0.24, χ^2^ = 1.66, *p* = 0.20, Figure 2). There was also no effect of cross-fostering timing (egg vs from P30) on overall song similarity (PC1) to tutor (ΔlogL = 0.83, χ^2^ = 0.48, *p* = 0.49). We did find, however, that birds tutored from the egg stage more closely matched their tutor in terms of pitch (ΔlogL = 2.82, χ^2^ = 5.64, *p* = 0.02) and log-transformed variance in entropy (ΔlogL = 3.38, χ^2^ = 6.75, *p* = 0.01), supporting a modest effect of cross-fostering approach on song similarity. Due to this effect, we use a balanced set of egg (n = 2) and P30 (n = 6) cross-fostered birds from the two subspecies to compare patterns song variability after cross-fostering. We found a significant effect of population on overall K-L distance (df = 1, χ^2^ = 18.47, *p* = 1.73e-05) and each individual song measurement (Figure 2). Thus, domesticated *T. g. castanotis* zebra finches maintained high song diversity than *T. g. guttata* even after cross-fostering.

**Figure 2.**
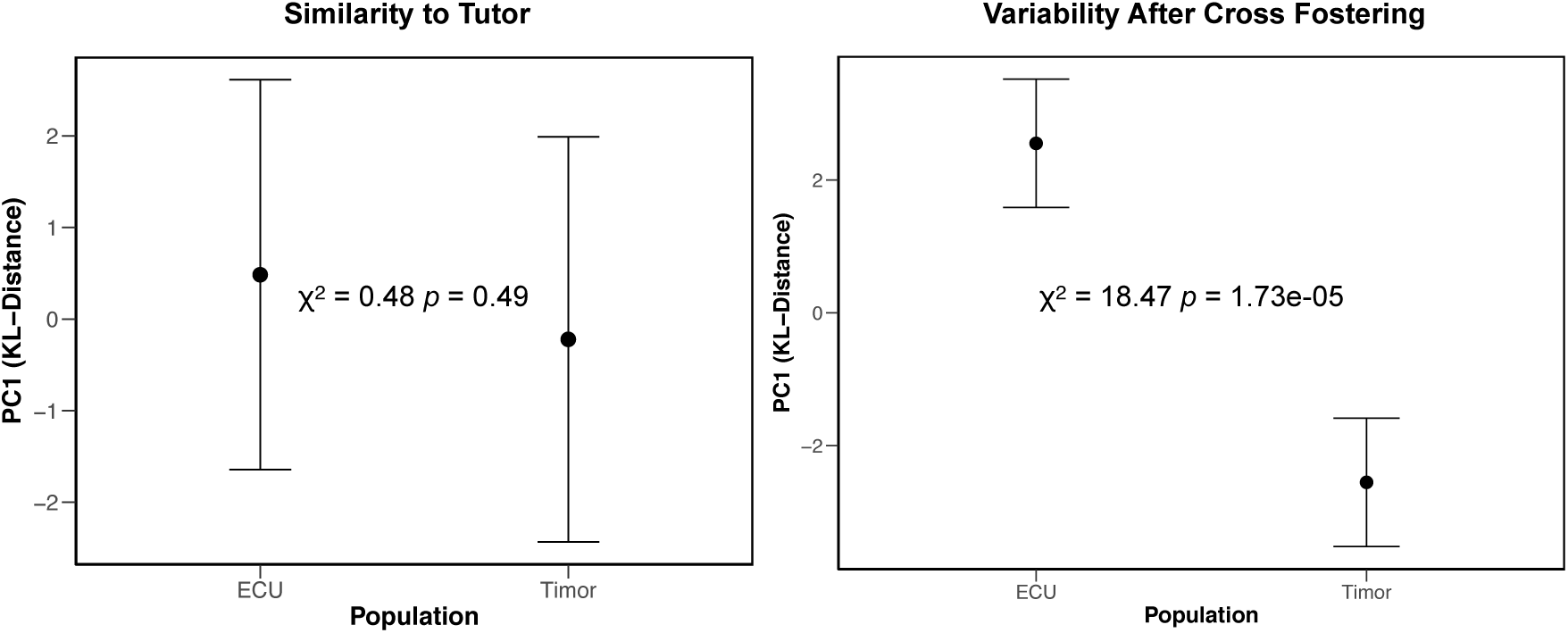
Fitted means and confidence intervals for K-L distance estimated A) between cross-fostered ECU and Timor birds and their respective tutors and B) within populations of cross-fostered birds. Estimated *χ*^2^ and associated p-value from likelihood ratio test are presented in both panels.

## 4. Discussion

Our analysis reveals differences in population-level song variability across zebra finch populations. We examined song variability in two domesticated populations of *T. g. castanotis* in the US and two more recently wild-derived populations, one of *T. g. castanotis* (Macquarie) and one of *T. g. guttata*. The two more recently wild-derived populations representing both subspecies had lower song variability than did the domesticated populations of *T. g. castanotis* in the US. While in principle, population differences in variability could be caused by either genetic or cultural evolution, our cross fostering experiment suggests that genetic differences contribute to this pattern, at least for the *T. g. guttata* and *T. g. castanotis* populations at ECU that were experimentally tested. These populations represent the extremes of song diversity in our survey. Following cross-fostering, domesticated *T. g. castanotis* maintained higher song variability than *T. g. guttata*. We suggest that higher song variability in domesticated *T. g. castanotis* could represent a consequence of reduced purifying selection on song characteristics in captivity. Previous studies of domesticated Bengalese finches have also shown high variability relative to the ancestral white-backed munia *Lonchura striata* (Okanoya 2004; Suzuki et al. 2014).

Although broadly consistent, high variability in domesticated zebra finch populations was not ubiquitous across populations or song parameters. The population sampled at the University of Chicago showed significantly higher song variability than wild-derived *T. g. castanotis* in only three out of thirteen song parameters, and not in overall K-L distance based on the summary principal component score. By contrast, the ECU population had higher mean K-L distance than each of the three other populations at every measured song parameter and the principal component summary measure. Aspects of population history therefore also play a role in shaping patterns of song variability.

By cross-fostering *T. g. castanotis* and *T. g. guttata* housed at ECU, we partially controlled for early developmental experience on song variability. The maintenance of high variability after cross fostering supports a genetic cause for the high variability in domesticated zebra finches. Three-quarters (6 out of 8) of the cross-fostered birds in each population, however, were tutored beginning at day 30 rather than from hatch. These birds would have been exposed to other birds from their population between P20 and P30, the earliest part of the sensory phase, and we cannot rule out that such exposure influenced subsequent song variability. Previous work in *T. g. castanotis* suggests that exposure to tutor song before P30 does not influence tutor song copying, whereas exposure beginning at P30 does (Roper and Zann 2006, London 2017). We did find, however, that birds tutored from the egg stage more closely matched their tutors in both pitch and variance in entropy.

There is increasing interest in the extent to which changes during domestication have impacted laboratory model systems. Domestication is known to influence aspects of social behavior such as reduced aggression and fear response, increased social cognitive abilities for interacting with humans, and reduced wild-type behavior towards humans (Frank and Frank 1982; Schütz et al. 2001; Hare et al. 2002; Trut et al. 2009). Recent studies have found that wild and domesticated zebra finches differ in an array of behaviors such as mate choice (Rustein et al. 2007), hatching synchrony (Mainwaring et al. 2010; Gilby et al. 2013), nest visit synchrony (Mariette and Griffith 2012), and parental care (Gilby et al. 2011). While there has been some documentation of laboratory populations of zebra finches exhibiting discrete vocal traditions from one another (Sturdy et al. 1999), the opposite has also been found (Lachlan et al. 2016). Our study reveals population differences in song variability itself. Recent work has begun to reveal genetic contributions to learned vocal communication behavior in songbirds (Mets & Brainard 2018) and song variability in zebra finches may offer a similar opportunity. As auditory experience shapes subsequent processing of acoustic stimuli (reviewed in Dong & Clayton 2009) consideration of these population differences will be important for future studies of vocal behavior and song recognition.

---

**Supplementary Figure 1**

Two-dimensional scatterplots of frequency against duration summarizing song repertoires for individual birds from the four study populations. Each point represents an individual syllable and clusters of points represents frequently repeated syllables. K-L distance measures the dissimilarity in the distribution of points between feature plots for pairs of birds.

## Ethical Statement

Animal use was conducted with approval from Institutional Animal Care and Use Committees of East Carolina University (AUP D285), the University of Chicago (ACUP72220). Animal use at Macquarie University was approved under AEC ARA 2015/017.

## Data Accessibility

R code and K-L distance measurements are available on GitHub (https://github.com/chrisbalakrishnan/SongVariability)

## Competing Interests

There are no competing interests.

## Author Contributions

ALL designed the study, conducted song recordings, carried out the experiments, analyzed the data and wrote the MS

CNB designed the study, conducted song recordings, analyzed the data and wrote the SEL conducted song recordings and edited the MS

DFC designed the study and edited the MS

SCG provided access to wild-derived birds and edited the MS

## Acknowledgements

We would like to thank Ken Soderstrom and Ofer Tchernichovsky for their guidance in this project and for extensive help with Sound Analysis Pro and K-L distance estimation. This work represents a portion of ALL’s dissertation project, which was greatly facilitated by committee members Ken Soderstrom, Carol Goodwillie, Michael Brewer, Jeff McKinnon and Sue McRae. Mike McCoy and Ariane Peralta provided guidance and help with code for the statistical analyses.

## Funding

This work was funded by East Carolina University and NIH NIGMS 1RC1GM091556-01 (DFC).

